## Supplemental Tables and Figures for "Development of D-box peptides to inhibit the Anaphase Promoting Complex/Cyclosome"

[b] AstraZeneca, Darwin building, 310 Milton Road, Cambridge, CB4 0FZ, UK.

[c] MRC Laboratory of Molecular Biology, Francis Crick Avenue, Cambridge, CB2 0QH, UK

[d] Yusuf Hamid Department of Chemistry, University of Cambridge, Lensfield Road, Cambridge, CB2 1EW, UK

[e] p53laboratory, 8A Biomedical Grove, #06-04/05, Neuros/Immunos, 138548, Singapore.

**Table S1.** Data collection, phasing and refinement statistics for Cdc20 crystal structures with D-box peptides

| Crystal structure | Cdc20 – D21 | Cdc20 – D20 | Cdc20 – D7 |
|---|---|---|---|
| PDB accession code | 9I68 | 9I69 | 9I6A |
| **Data collection** | | | |
| Space group | $P2_1$ | $P2_1$ | $P2_1$ |
| Unit cell, a, b, c (Å),<br>α, β, γ (°) | 35.49, 87.55, 48.57<br>90.00, 109.71, 90.00 | 35.34, 87.34, 48.51<br>90.00, 110.14, 90.00 | 35.00, 86.87, 48.03<br>90.00, 109.60, 90.00 |
| Resolution range, Å | 45.73 - 1.51 (1.66 - 1.51) | 45.55 - 1.46 (1.60 - 1.46) | 45.24 - 1.92 (2.09 - 1.92) |
| Total reflections | 162715 (6076) | 184416 (10465) | 76640 (3657) |
| Unique reflections | 32820 (1641) | 35450 (1772) | 14979 (750) |
| Multiplicity | 5.0 (3.7) | 5.2 (5.9) | 5.1 (4.9) |
| Completeness (spherical), % | 74.7 (14.5) | 74.5 (16.0) | 71.8 (15.4) |
| Completeness (ellipsoidal), % | 90.4 (41.9) | 92.9 (60.1) | 90.5 (57.6) |
| I/σI | 13.1 (1.6) | 13.6 (1.4) | 7.6 (1.5) |
| $R_{merge}$ | 0.050 (0.598) | 0.048 (1.000) | 0.132 (1.161) |
| $CC_{1/2}$ | 0.999 (0.714) | 0.999 (0.635) | 0.996 (0.536) |
| **Refinement** | | | |
| $R_{work}/R_{free}$, % | 0.175/0.194 | 0.165/0.184 | 0.207/0.231 |
| Unique reflections used | 32820 | 35450 | 14966 |
| rmsd bond lengths, Å | 0.008 | 0.008 | 0.008 |
| rmsd bond angles, ° | 1.01 | 1.02 | 1.00 |
| Ramachandran analysis: | | | |
| Favoured, % | 97.10 | 98.06 | 97.33 |
| Allowed, % | 2.58 | 1.61 | 2.33 |
| Outliers, % | 0.32 | 0.32 | 0.33 |
| Number of atoms<br>(average B-factor, Å$^2$) | | | |
| Protein | 2452 (24.31) | 2434 (25.80) | 2340 (29.68) |
| Peptide | 46 (37.72) | 46 (46.88) | 25 (48.49) |
| Solvent | 236 (39.17) | 240 (44.08) | 72 (32.96) |
| Mean/Wilson B-factor, Å$^2$ | 25.8/22.9 | 27.8/23.9 | 30.0/27.8 |

**Figure S1.**

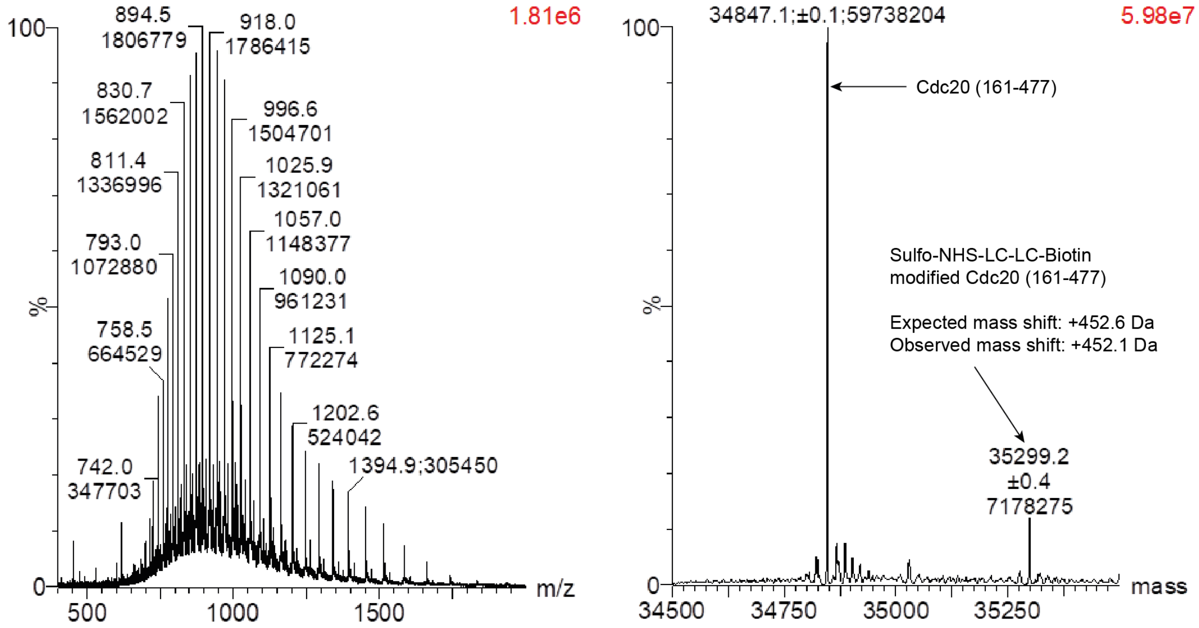


**Figure S1.** Verification of single-site biotinylation of purified Cdc20 WD40 domain (residues 161-477) using Sulfo-NHS-LC-LC-Biotin (Thermo Scientific, A35358). The observed mass shift corresponds to one Sulfo-NHS-LC-LC-Biotin molecule added to the purified protein.

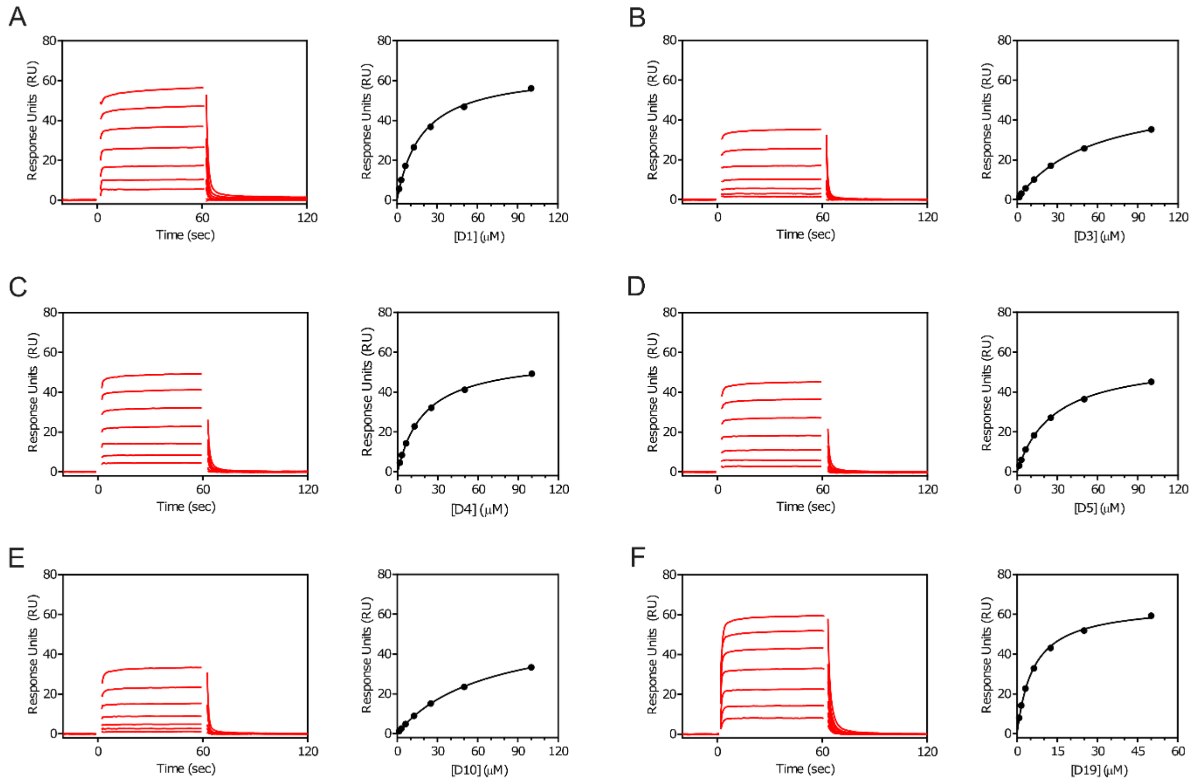


**Figure S2.** SPR reference-subtracted sensorgrams and binding curves for **(A)** D1, **(B)** D3, **(C)** D4, **(D)** D5, **(E)** D10, **(F)** D19.

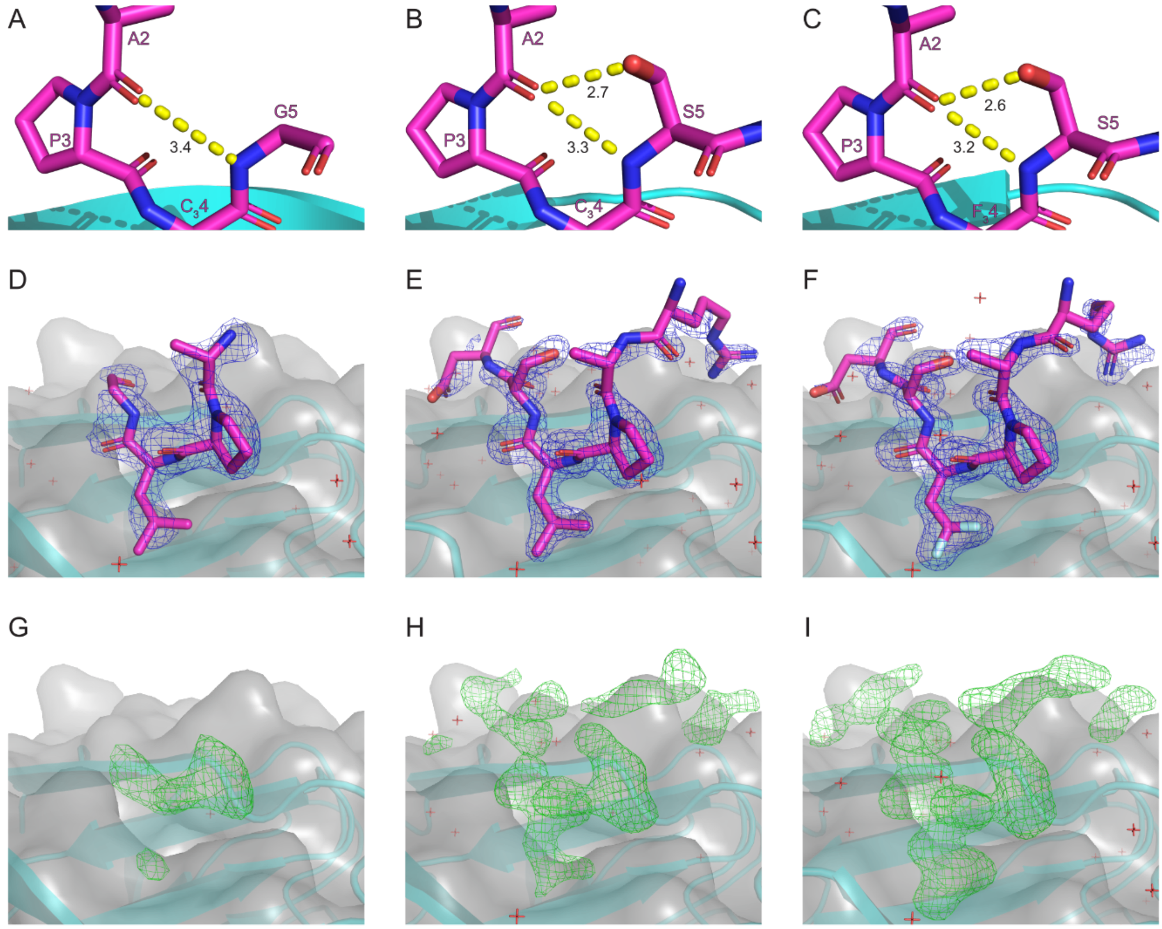


**Figure S3**. Cdc20$^{WD40}$ is shown in cartoon (cyan) and surface (grey) representations. All peptides are shown by stick representations (magenta). Intramolecular hydrogen bonds formed within peptides **(A)** D7, **(B)**, D20 and **(C)** D21. All peptides form a hydrogen bond between the carbonyl of A2 to the amine of G5/S5. Peptides D20 and D21 form an additional H-bond between the carbonyl of A2 to the hydroxyl group of S5. $2F_oF_c$ maps contoured at 1.0σ and modelled peptide atoms of **(D)** D7, **(E)**, D20 and **(F)** D21. Unbiased $F_oF_c$ maps from refinement steps prior to including the peptides in the model, contoured at 2.5σ. Maps shown are, **(G)** D7, **(H)** D20 and **(I)** D21.

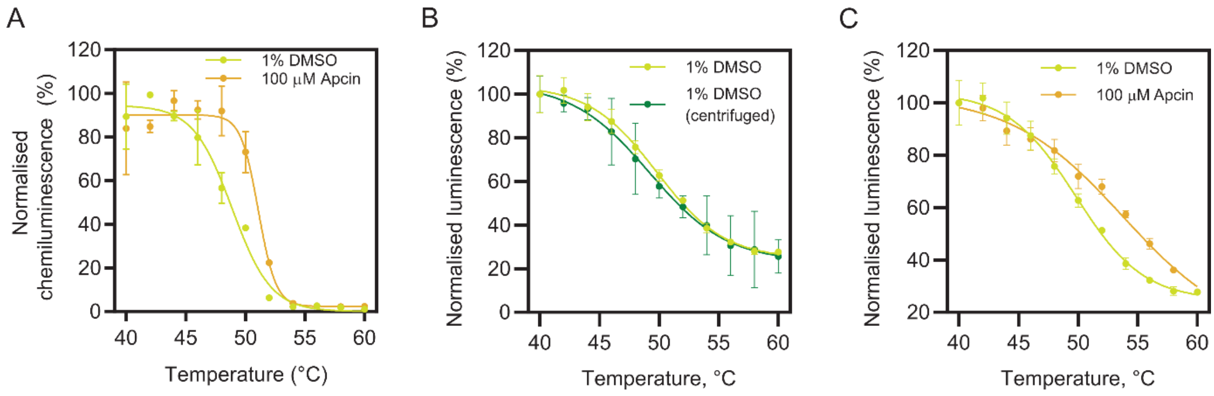


**Figure S4**. (A) CETSA of endogenous Cdc20 in HEK293T cell lysates by 100 µM Apcin compared with vehicle control (1% DMSO), analysed by densitometric of western blots. Mean and standard deviation are calculated from two independent experiments. (B) CETSA of transfected Cdc20 with a C-terminal HiBiT tag spiked with 1% DMSO with and without including a centrifugation step following heat denaturation at each temperature set point. (C) Stabilisation of transfected Cdc20 with a C-terminal HiBiT tag by 100 µM Apcin compared with vehicle control (1% DMSO).

#### D1: Ac-GRAALSDITN-$NH_2$

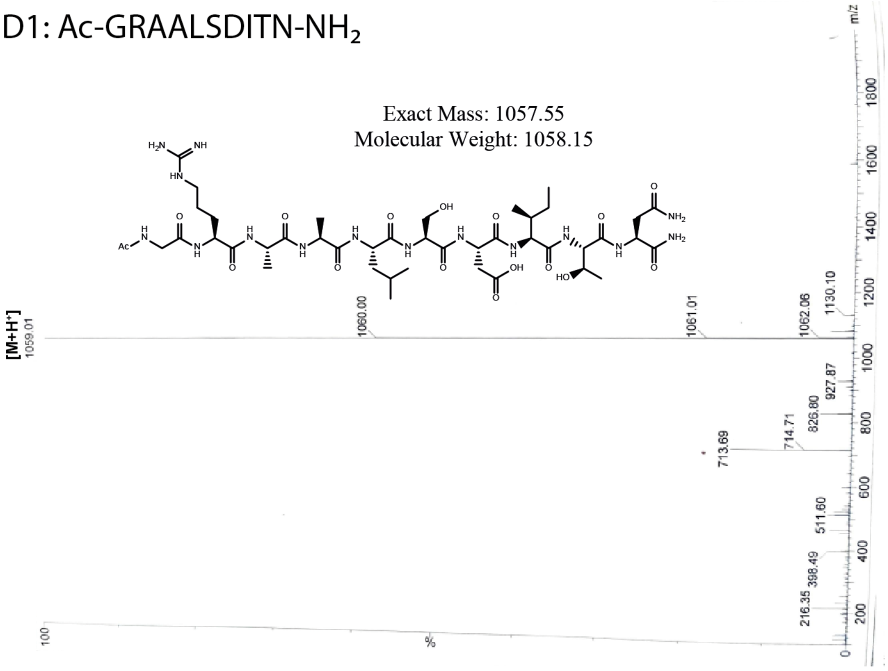


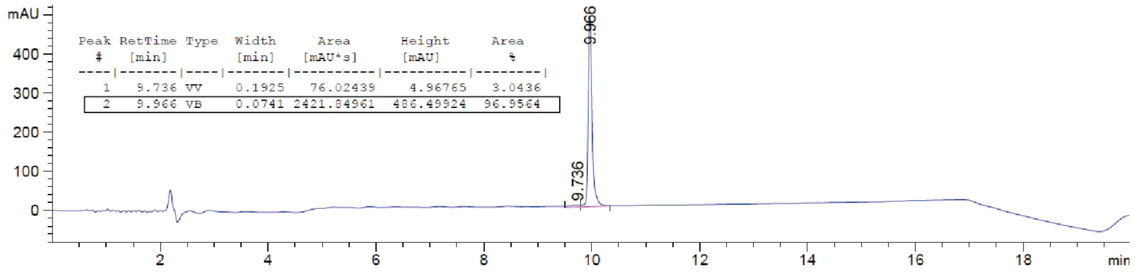


| Peak # | RetTime [min] | Type | Width [min] | Area [mAU*s] | Height [mAU] | Area % |
|---|---|---|---|---|---|---|
| 1 | 9.736 | VV | 0.1925 | 76.02439 | 4.96765 | 3.0436 |
| 2 | 9.966 | VB | 0.0741 | 2421.84961 | 486.49924 | 96.9564 |

#### D2: Ac-RLPLGDVSN-$NH_2$

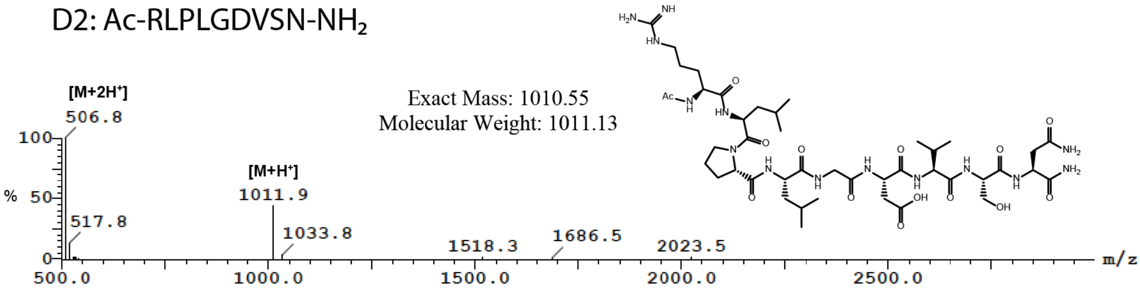


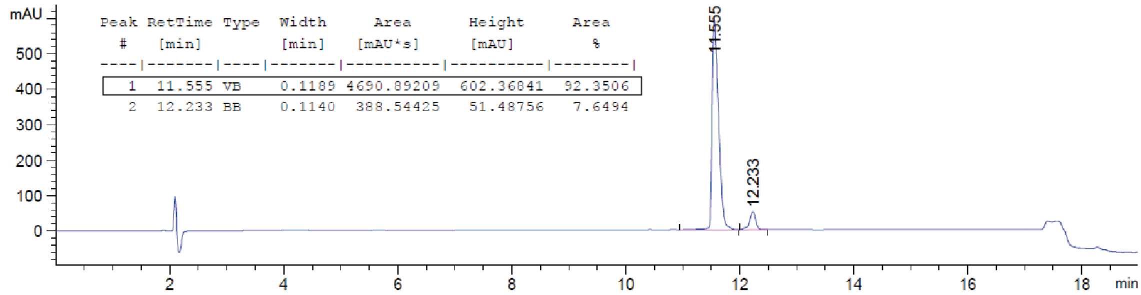


| Peak # | RetTime [min] | Type | Width [min] | Area [mAU*s] | Height [mAU] | Area % |
|---|---|---|---|---|---|---|
| 1 | 11.555 | VB | 0.1189 | 4690.89209 | 602.36841 | 92.3506 |
| 2 | 12.233 | BB | 0.1140 | 388.54425 | 51.48756 | 7.6494 |

### D3: Ac-RAPLGDVSN-$NH_2$

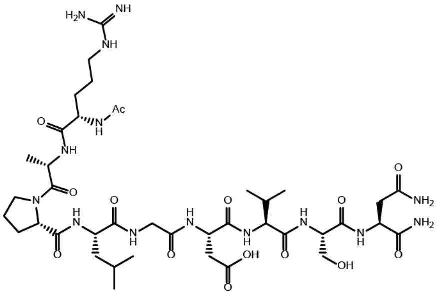

Exact Mass: 968.50
Molecular Weight: 969.05

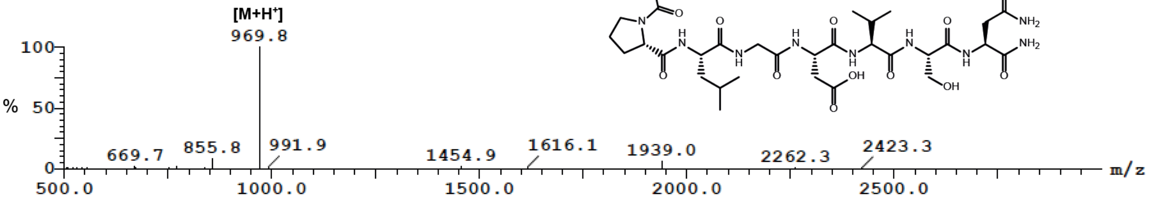


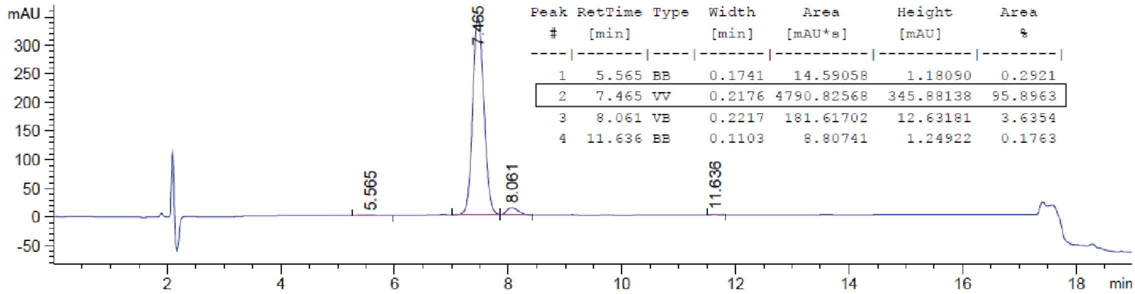


| Peak # | RetTime [min] | Type | Width [min] | Area [mAU*s] | Height [mAU] | Area % |
|---|---|---|---|---|---|---|
| 1 | 5.565 | BB | 0.1741 | 14.59058 | 1.18090 | 0.2921 |
| 2 | 7.465 | VV | 0.2176 | 4790.82568 | 345.88138 | 95.8963 |
| 3 | 8.061 | VB | 0.2217 | 181.61702 | 12.63181 | 3.6354 |
| 4 | 11.636 | BB | 0.1103 | 8.80741 | 1.24922 | 0.1763 |

### D4: Ac-RAPLGDISN-$NH_2$

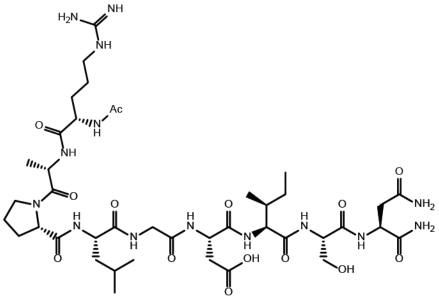

Exact Mass: 982.52
Molecular Weight: 983.08

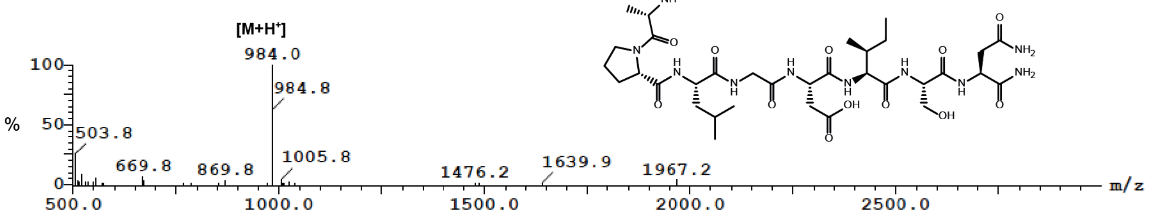


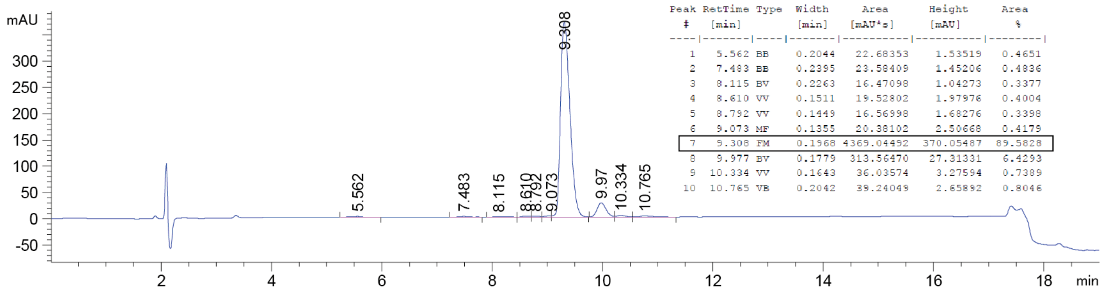


| Peak # | RetTime [min] | Type | Width [min] | Area [mAU*s] | Height [mAU] | Area % |
|---|---|---|---|---|---|---|
| 1 | 5.562 | BB | 0.2044 | 22.68353 | 1.53519 | 0.4651 |
| 2 | 7.483 | BB | 0.2395 | 23.58409 | 1.45206 | 0.4836 |
| 3 | 8.115 | BV | 0.2263 | 16.47098 | 1.04273 | 0.3377 |
| 4 | 8.610 | VV | 0.1511 | 19.52802 | 1.97976 | 0.4004 |
| 5 | 8.792 | VV | 0.1449 | 16.56998 | 1.68276 | 0.3398 |
| 6 | 9.073 | MF | 0.1355 | 20.38102 | 2.50668 | 0.4179 |
| 7 | 9.308 | FM | 0.1968 | 4369.04492 | 370.05487 | 89.5828 |
| 8 | 9.977 | BV | 0.1779 | 313.56470 | 27.31331 | 6.4293 |
| 9 | 10.334 | VV | 0.1643 | 36.03574 | 3.27594 | 0.7389 |
| 10 | 10.765 | VB | 0.2042 | 39.24049 | 2.65892 | 0.8046 |

#### D5: Ac-RAPLGDLSN-$NH_2$

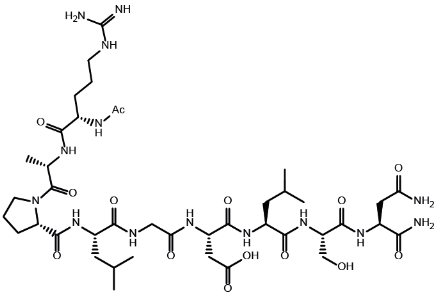

Exact Mass: 982.52
Molecular Weight: 983.08

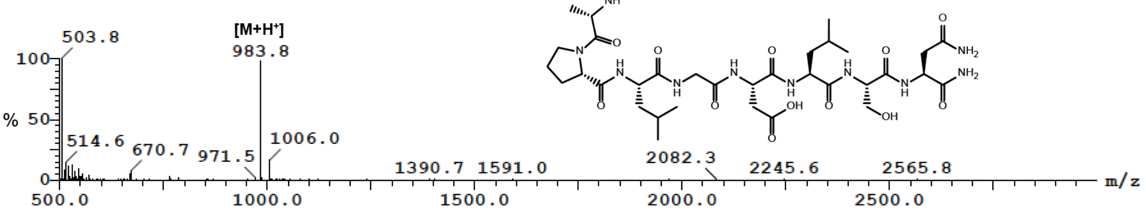


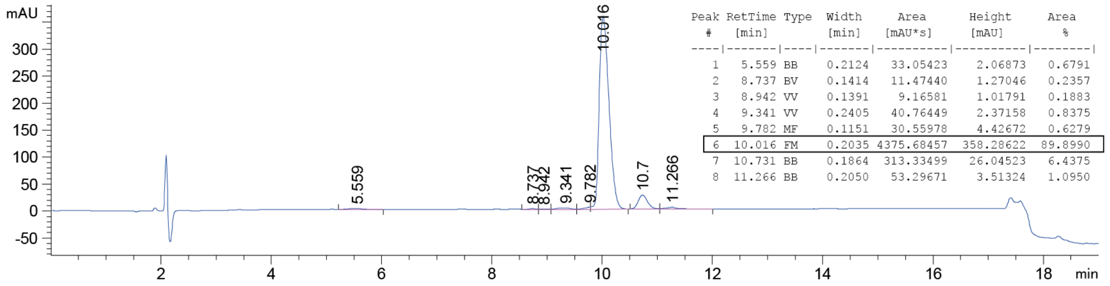


| Peak # | RetTime [min] | Type | Width [min] | Area [mAU*s] | Height [mAU] | Area % |
|---|---|---|---|---|---|---|
| 1 | 5.559 | BB | 0.2124 | 33.05423 | 2.06873 | 0.6791 |
| 2 | 8.737 | BV | 0.1414 | 11.47440 | 1.27046 | 0.2357 |
| 3 | 8.942 | VV | 0.1391 | 9.16581 | 1.01791 | 0.1883 |
| 4 | 9.341 | VV | 0.2405 | 40.76449 | 2.37158 | 0.8375 |
| 5 | 9.782 | MF | 0.1151 | 30.55978 | 4.42672 | 0.6279 |
| 6 | 10.016 | FM | 0.2035 | 4375.68457 | 358.28622 | 89.8990 |
| 7 | 10.731 | BB | 0.1864 | 313.33499 | 26.04523 | 6.4375 |
| 8 | 11.266 | BB | 0.2050 | 53.29671 | 3.51324 | 1.0950 |

#### D7: Ac-RAP**$C_3$**GDISN-$NH_2$

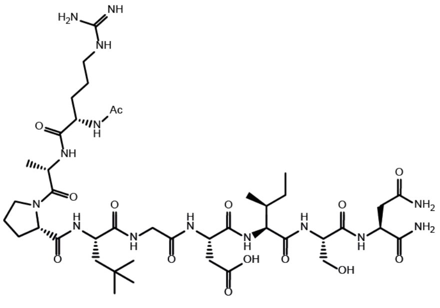

Exact Mass: 996.54
Molecular Weight: 997.11

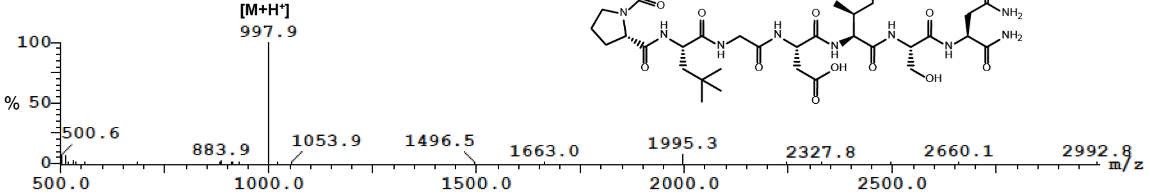


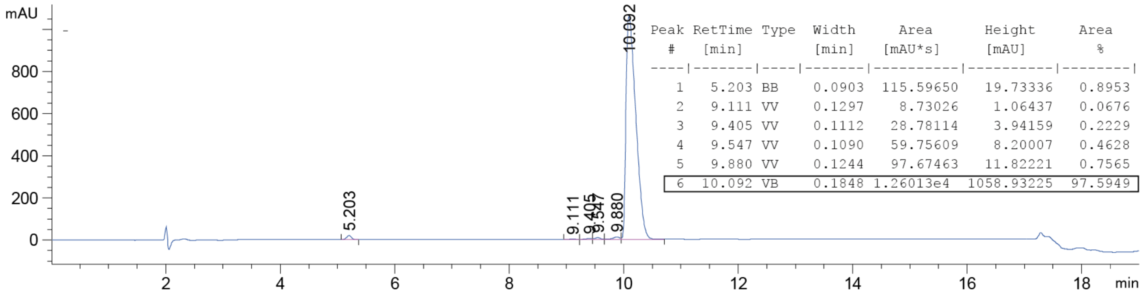


| Peak # | RetTime [min] | Type | Width [min] | Area [mAU*s] | Height [mAU] | Area % |
|---|---|---|---|---|---|---|
| 1 | 5.203 | BB | 0.0903 | 115.59650 | 19.73336 | 0.8953 |
| 2 | 9.111 | VV | 0.1297 | 8.73026 | 1.06437 | 0.0676 |
| 3 | 9.405 | VV | 0.1112 | 28.78114 | 3.94159 | 0.2229 |
| 4 | 9.547 | VV | 0.1090 | 59.75609 | 8.20007 | 0.4628 |
| 5 | 9.880 | VV | 0.1244 | 97.67463 | 11.82221 | 0.7565 |
| 6 | 10.092 | VB | 0.1848 | 1.26013e4 | 1058.93225 | 97.5949 |

#### D10: Ac-RAALGDISN-$NH_2$

Exact Mass: 956.50
Molecular Weight: 957.04

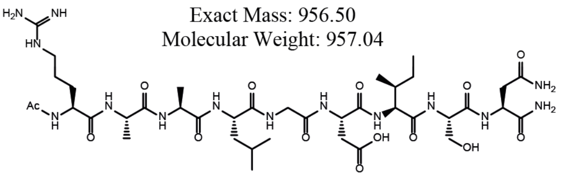

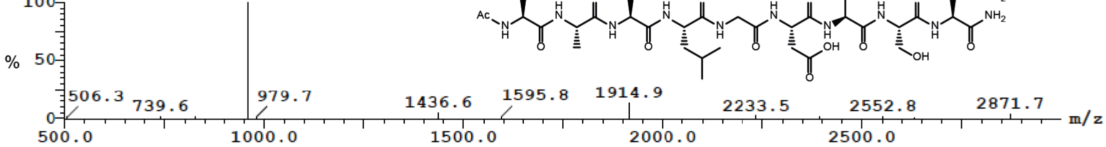


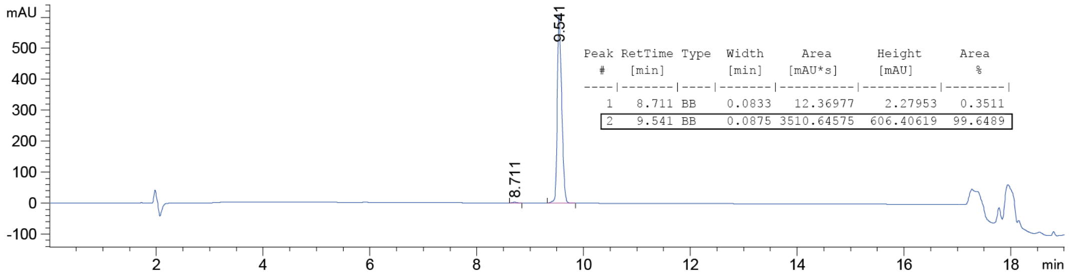


| Peak # | RetTime [min] | Type | Width [min] | Area [mAU*s] | Height [mAU] | Area % |
|---|---|---|---|---|---|---|
| 1 | 8.711 | BB | 0.0833 | 12.36977 | 2.27953 | 0.3511 |
| 2 | 9.541 | BB | 0.0875 | 3510.64575 | 606.40619 | 99.6489 |

#### D12: Ac-RAA**C₃**GDISN-$NH_2$

Exact Mass: 970.52
Molecular Weight: 971.07

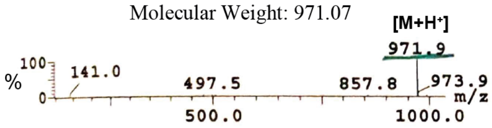


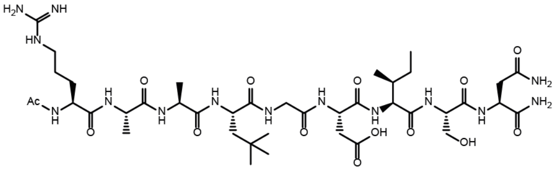

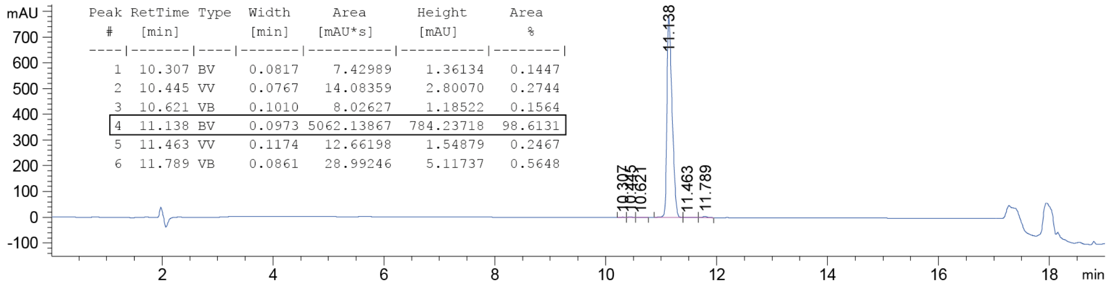


| Peak # | RetTime [min] | Type | Width [min] | Area [mAU*s] | Height [mAU] | Area % |
|---|---|---|---|---|---|---|
| 1 | 10.307 | BV | 0.0817 | 7.42989 | 1.36134 | 0.1447 |
| 2 | 10.445 | VV | 0.0767 | 14.08359 | 2.80070 | 0.2744 |
| 3 | 10.621 | VB | 0.1010 | 8.02627 | 1.18522 | 0.1564 |
| 4 | 11.138 | BV | 0.0973 | 5062.13867 | 784.23718 | 98.6131 |
| 5 | 11.463 | VV | 0.1174 | 12.66198 | 1.54879 | 0.2467 |
| 6 | 11.789 | VB | 0.0861 | 28.99246 | 5.11737 | 0.5648 |

#### D19: Ac-RAPLSDITN-$NH_2$

Exact Mass: 1026.55
Molecular Weight: 1027.13

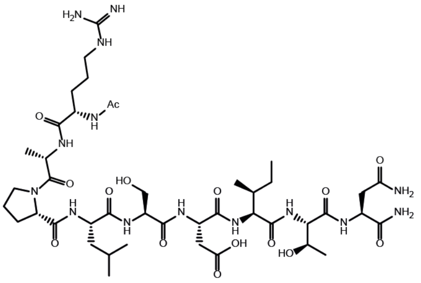

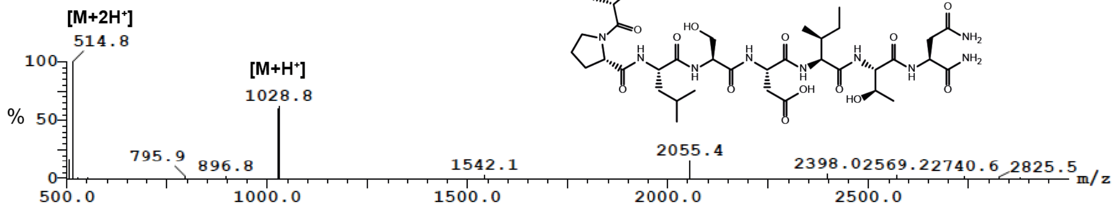


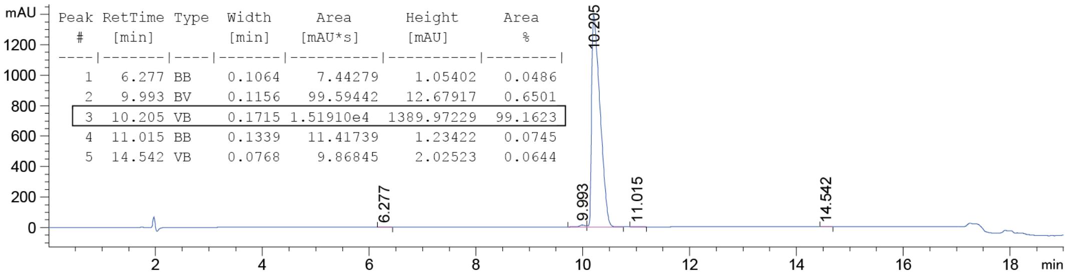


| Peak # | RetTime [min] | Type | Width [min] | Area [mAU*s] | Height [mAU] | Area % |
|---|---|---|---|---|---|---|
| 1 | 6.277 | BB | 0.1064 | 7.44279 | 1.05402 | 0.0486 |
| 2 | 9.993 | BV | 0.1156 | 99.59442 | 12.67917 | 0.6501 |
| 3 | 10.205 | VB | 0.1715 | 1.51910e4 | 1389.97229 | 99.1623 |
| 4 | 11.015 | BB | 0.1339 | 11.41739 | 1.23422 | 0.0745 |
| 5 | 14.542 | VB | 0.0768 | 9.86845 | 2.02523 | 0.0644 |

D20: Ac-RAP**C₃**SDITN-NH₂

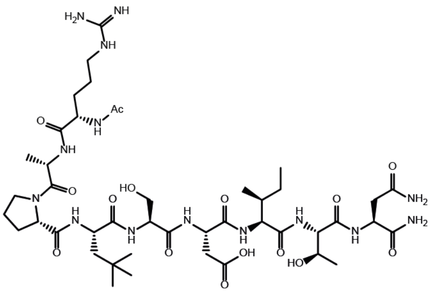

Exact Mass: 1040.56
Molecular Weight: 1041.16

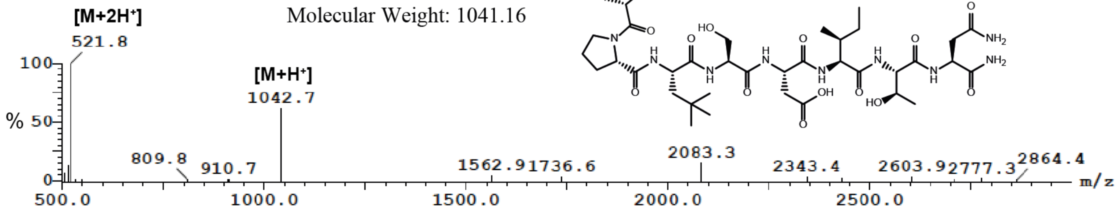


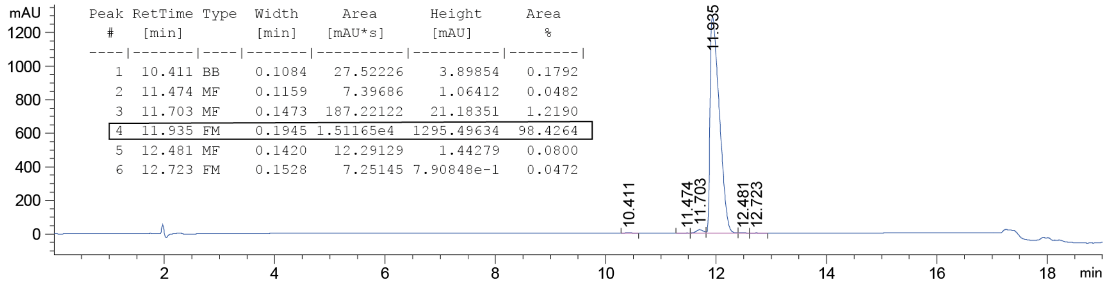


| Peak # | RetTime [min] | Type | Width [min] | Area [mAU*s] | Height [mAU] | Area % |
|---|---|---|---|---|---|---|
| 1 | 10.411 | BB | 0.1084 | 27.52226 | 3.89854 | 0.1792 |
| 2 | 11.474 | MF | 0.1159 | 7.39686 | 1.06412 | 0.0482 |
| 3 | 11.703 | MF | 0.1473 | 187.22122 | 21.18351 | 1.2190 |
| 4 | 11.935 | FM | 0.1945 | 1.51165e4 | 1295.49634 | 98.4264 |
| 5 | 12.481 | MF | 0.1420 | 12.29129 | 1.44279 | 0.0800 |
| 6 | 12.723 | FM | 0.1528 | 7.25145 | 7.90848e-1 | 0.0472 |

D21: Ac-RAP**F₃**SDITN-NH₂

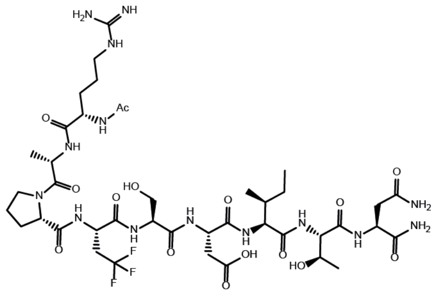

Exact Mass: 1052.49
Molecular Weight: 1053.05

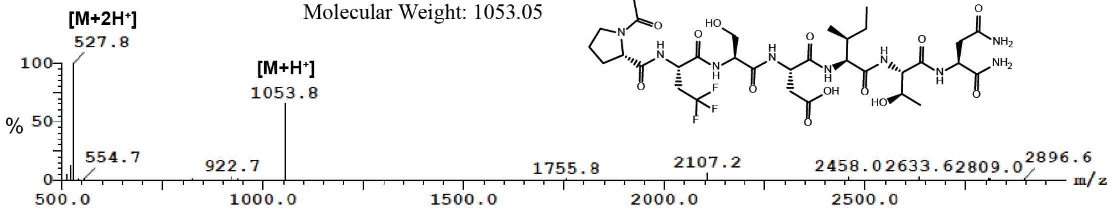


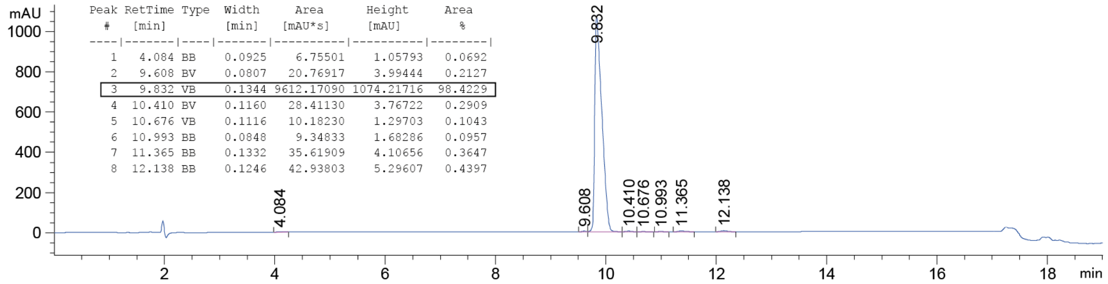


| Peak # | RetTime [min] | Type | Width [min] | Area [mAU*s] | Height [mAU] | Area % |
|---|---|---|---|---|---|---|
| 1 | 4.084 | BB | 0.0925 | 6.75501 | 1.05793 | 0.0692 |
| 2 | 9.608 | BV | 0.0807 | 20.76917 | 3.99444 | 0.2127 |
| 3 | 9.832 | VB | 0.1344 | 9612.17090 | 1074.21716 | 98.4229 |
| 4 | 10.410 | BV | 0.1160 | 28.41130 | 3.76722 | 0.2909 |
| 5 | 10.676 | VB | 0.1116 | 10.18230 | 1.29703 | 0.1043 |
| 6 | 10.993 | BB | 0.0848 | 9.34833 | 1.68286 | 0.0957 |
| 7 | 11.365 | BB | 0.1332 | 35.61909 | 4.10656 | 0.3647 |
| 8 | 12.138 | BB | 0.1246 | 42.93803 | 5.29607 | 0.4397 |

**Figure S5.** Identity and quality control information for all peptides synthesised and purified in this study. Linear amino acid sequence and chemical structures are illustrated alongside the expected exact mass and molecular weights for each peptide. LCMS data showing the *m/z* species for each purified peptide. Analytical HPLC chromatograms (10-30% solvent B over 15 mins, 1 mL/min, UV trace at 220 nm), with calculated purity of the peptides by the percentage area.
